# Computational Redesign of an Antifreeze Protein Using Deep Learning

**DOI:** 10.64898/2026.06.21.733612

**Authors:** Cianna Calia, Arthur J. Altunc, Rosemary J. Eufemio, Benjamin O. Alvarado, Jake D. Huynh, Eunkeu Oh, Michael D. Burkart, Konrad Meister, Francesco Paesani

## Abstract

Antifreeze proteins (AFPs) found in various cold-adapted organisms inhibit ice growth and are of interest for applications in food products, cryopreservation, agriculture, and materials science. Although high-resolution structures are available for several AFPs, the amino acids required for full antifreeze activity remain incompletely defined, and the development of AFP variants with properties such as enhanced solubility, high expression yield, and improved thermostability may further facilitate applications. Here, we used the deep learning model ProteinMPNN to redesign the globular fish antifreeze protein AFPIII, keeping the previously reported ice-binding residues fixed. We readily obtained sequences confidently predicted to adopt AFPIII’s structure and we selected five designed variants for expression, all of which expressed efficiently in *E. coli*. Circular dichroism spectroscopy showed that two of these variants retained secondary structure elements consistent with AFPIII, whereas the other three exhibited structural differences. One design was predicted and experimentally confirmed to have increased thermostability. All five variants displayed measurable thermal hysteresis activity. However, none reached the activity of wild-type AFPIII, suggesting that maintaining the currently established set of ice-binding residues is not sufficient to fully preserve this AFP’s function; other, unidentified residues can also impact its activity. Our findings highlight the value of deep learning-based protein design methods both for generating AFP variants with desirable properties and for uncovering gaps in existing knowledge of well-characterized AFPs.

## Introduction

Deep learning methods for protein structure prediction and design have advanced rapidly in recent years (e.g., Refs. 1–16). Generative design models coupled with structure prediction tools now enable the production and computational screening of designed proteins under a variety of defined constraints. These approaches have created highly active enzymes, ^17–23^ de novo binders for therapeutic targets,^10,15,24–31^ biosensors for small molecules, ^32–34^ and multi-component assemblies. ^35–39^ The use of such methods to develop new biomolecules that interact with and modulate the growth of inorganic crystals presents another exciting frontier.^40–43^ In addition to de novo design, deep learning enables systematic modification of existing natural proteins to probe structure-function relationships while improving application-relevant properties such as expression yield, stability, and tolerance to environmental stress. ^17^ Natural protein classes with high translational potential but limited current usability therefore represent attractive targets for computational redesign.

A range of cold-adapted organisms, including fish, insects, plants, and microorganisms, express ice-binding proteins (IBPs) that facilitate their survival in icy habitats.^44–47^ Antifreeze proteins (AFPs) are a well-characterized subset of IBPs that inhibit macroscopic ice growth through adsorption to specific crystallographic planes. Upon adsorption to ice, AFPs restrict ice growth to regions between adsorbed proteins, generating interfacial curvature. This manifestation of the Kelvin effect results in thermal hysteresis, defined as the gap between the nonequilibrium freezing temperature and the melting temperature.^48–51^ Many AFPs also inhibit ice recrystallization, the process in which larger ice crystals grow at the expense of smaller ones. ^52,53^

AFP structures found in nature range from beta-solenoid structures in insect AFPs^54,55^ to a single alpha-helix in the winter flounder type I AFP^56^ to a compact globular fold for the 7-kDa type III AFP from eelpouts (AFPIII).^57,58^ Despite the structural differences, all AFPs possess a dedicated ice-binding site (IBS) responsible for their interaction with ice, frequently mediated by preordered interfacial water molecules. ^55,59–74^ These sites are typically large, flat, slightly hydrophobic surfaces, often with repetitive elements. For example, the repeating threonine-X-threonine motifs found in insect AFPs bind to the basal and prism planes of ice.^63^ In contrast, AFPIII contains a nonrepetitive IBS that interacts with the pyramidal and primary prism planes; mutational studies have identified residues within this IBS that are required for thermal hysteresis activity. ^59,75–80^

Besides providing a scaffold for the IBS, the functional significance of the rest of an AFP’s structure and surface (the non-IBS) is only beginning to be understood. Molecular dynamics (MD) simulations of AFPs in solution have suggested that non-ice-binding surfaces of AFPs have a disordering impact on nearby waters, potentially helping bound AFPs resist engulfment by ice.^65,81^ The distinct behavior of the IBS versus the non-IBS is further supported by the observation of opposing effects on ice nucleation depending on the orientation of AFPs tethered to surfaces.^82^ It has been noted that AFPs frequently possess exposed charged residues in their non-IBS, ^83^ but simulations of the solenoidal spruce budworm AFP bound to ice and a mutant with all non-IBS charged residues replaced with alanine suggested that the overall shape of the non-IBS may be more critical than its surface chemistry for resisting engulfment by ice.^84^ New experimental comparisons that substantially alter AFPs’ non-IBS surface chemistry while minimizing changes in structure are therefore of interest for the continued study of engulfment resistance mechanisms.

AFPs have seen their first real-world application in frozen food, ^85–87^ and their potential is also being investigated for uses in cryopreservation, ^88–93^ frost-resistant crops, ^94–97^ and anti-icing materials.^98–103^ For these applications, expression yield, solubility, and structural stability are important in addition to activity. Natural AFPs do not necessarily optimize these properties simultaneously;^104^ combined with high production costs, this explains the limited translation of AFPs’ potential to actual applications.

The demand for active, readily usable AFPs has prompted efforts to diversify and optimize natural AFPs. AFP engineering and optimization strategies to date have included structure-informed residue substitutions,^105,106^ extension of repetitive AFP structures to increase the area of the IBS,^107,108^ surface modification through chemical cationization, ^109^ and optimization of simulated properties using a genetic algorithm.^110^ The recent emergence of many deep learning methods for protein design presents a variety of new possibilities to design and engineer IBPs, and these tools are only beginning to be applied in the context of ice growth modulation.^111,112^

Here we employed the deep learning model ProteinMPNN^9^ to redesign AFPIII while pre-serving its established ice-binding residues. ProteinMPNN generates amino acid sequences for a specified three-dimensional backbone structure and has been shown to produce designs with high stability and expression efficiency.^17^ Our primary aims were to develop AFPIII variants with improved practical usability and to determine how extensive sequence modifications outside the established IBS affect thermal hysteresis activity. After screening sequences with structure prediction tools, we selected five AFPIII variants for expression and characterization, all with substantial changes in their non-IBS regions.

## Results

### Generation of alternative AFPIII sequences with ProteinMPNN

We selected PDB 1HG7^78^ as a starting point for the structure-conditioned generation of alternative AFPIII sequences, as outlined in Fig. 1a. This crystal structure represents a recombinant version of AFPIII called rQAE m1.1 that has slight alterations from the ocean pout (*Macrozoarces americanus*) sequence but has been regarded as representative of the natural protein in prior studies. ^58,59^ We thus refer to rQAE m1.1 as wild-type AFPIII (WT AFPIII) in this manuscript and number the residues based on positions in the natural sequence, which lacks the initial methionine. To produce additional starting structures, we obtained five predicted structures for this sequence using the ColabFold implementation^113^ of AlphaFold 2^1^ (AF2).

**Figure 1:**
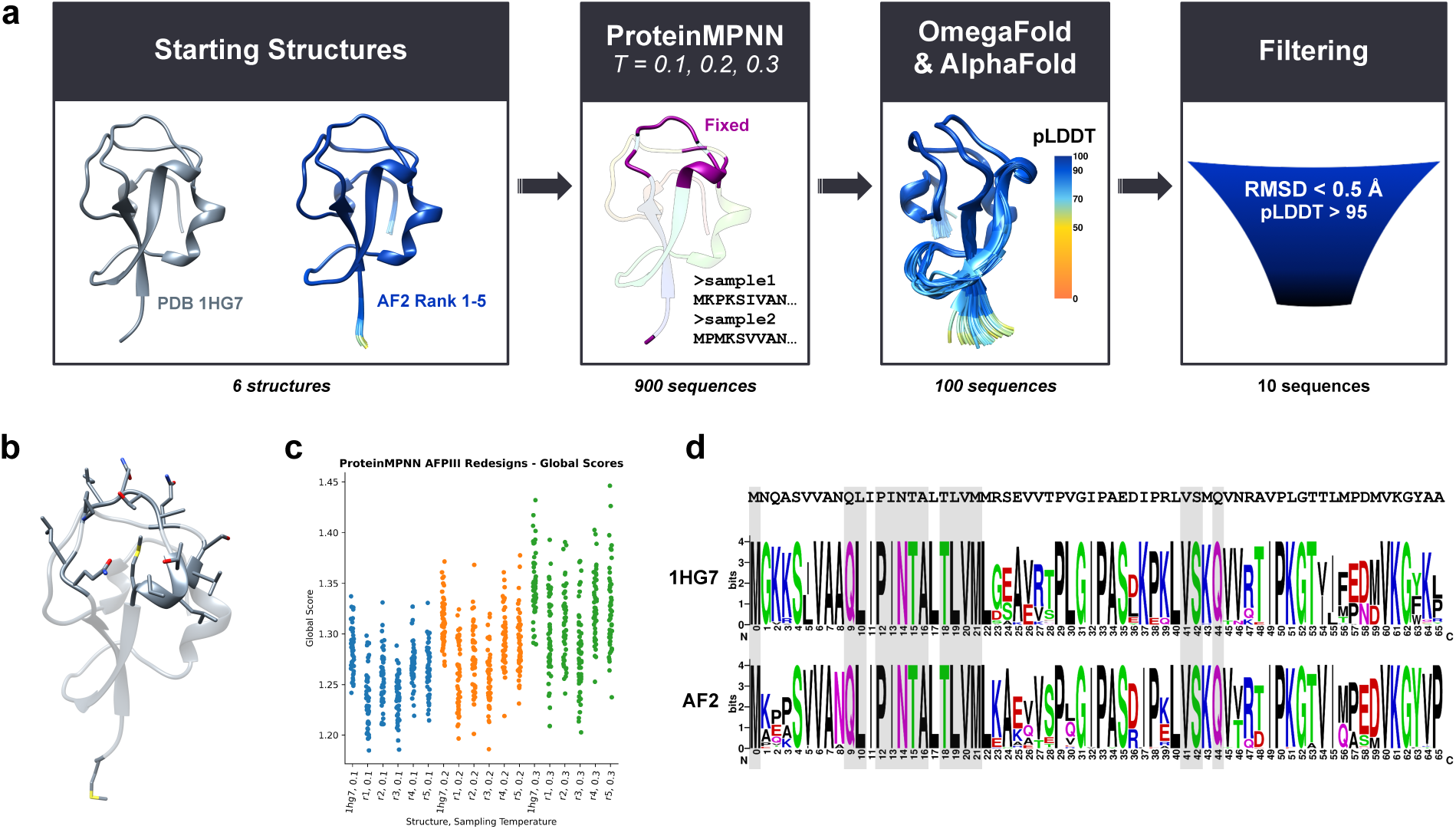
(a) Overview of our workflow for generating and screening alternative AFPIII sequences. (b) Crystal structure of AFPIII (PDB 1HG7) with side chains shown for the residues that we kept fixed in the redesign process. (c) Global scores from ProteinMPNN (average negative log likelihood over all residues given the corresponding structure) for all AFPIII variant sequences generated. For comparison, the global score of WT AFPIII’s sequence was 1.6920 for the crystal structure and ranged from 1.6026 to 1.6724 for the AlphaFold structures. (d) Sequence logos representing the 50 sequences generated based on PDB 1HG7 (top logo) and the 50 based on AlphaFold structures (bottom logo) that were selected for further analysis. The WT sequence is displayed above the logos. Fixed positions are shaded in gray.

Each of the six structures (PDB 1HG7 and the predicted structures) was provided as input to ProteinMPNN, with which we generated 150 cysteine-free sequences per structure, split between three sampling temperatures. We fixed the identity of residues in the protein’s compound IBS and also the N-terminal methionine (Fig. 1b). The fixed IBS amino acids included all IBS residues identified by Garnham et al.,^80^ as well as two other exposed residues (Pro12 and Ser42) that lie in the same planes.

The distribution of ProteinMPNN’s global score values (the average negative log probability over all residues given the corresponding backbone) differed noticeably between sequences generated based on the crystal structure and those based on the predicted structures, with scores for the former tending to be higher (Fig. 1c). We selected 100 sequences for further screening: the 50 sequences with the best (lowest) global scores among those from AF2 structures and the 50 best-scoring sequences from PDB 1HG7. These two sets exhibited substantial distinctions in amino acid frequencies at certain positions (Fig. 1d), illustrating that our use of multiple very similar structures as input to ProteinMPNN had contributed to the diversity of the designs. For example, the sequences from AF2 structures predominantly preserved the identity of the IBS-adjacent residue Asn8, unlike the designs from PDB 1HG7 in which this residue was consistently changed to alanine.

### In silico filtering of designed sequences with structure prediction models

We employed two different structure prediction models, OmegaFold (OF)^4^ and AF2, to identify the designs most confidently predicted to adopt the intended AFPIII fold. Structures of all 100 selected sequences were first predicted using OF. All designs were predicted to preserve AFPIII’s fold with fairly high to very high confidence (Fig. 2a), with the lowest average pLDDT (predicted local distance difference test) value being 85.7. Structural variability was primarily located near the termini and in a loop opposite the IBS (indicated by the arrow in Fig. 2a). Only sequences with an average OF pLDDT greater than 95 and an alpha carbon root mean squared deviation (RMSD) below 0.5 Å relative to PDB 1HG7 were kept for further analysis. To account for expected intrinsic flexibility near the termini, the first and last two residues’ alpha carbon atoms were excluded from the RMSD calculations. Of the 15 designs meeting these criteria, 13 had been generated using AF2-derived structures of wild-type AFPIII as input backbones, whereas two were based on the crystal structure.

**Figure 2:**
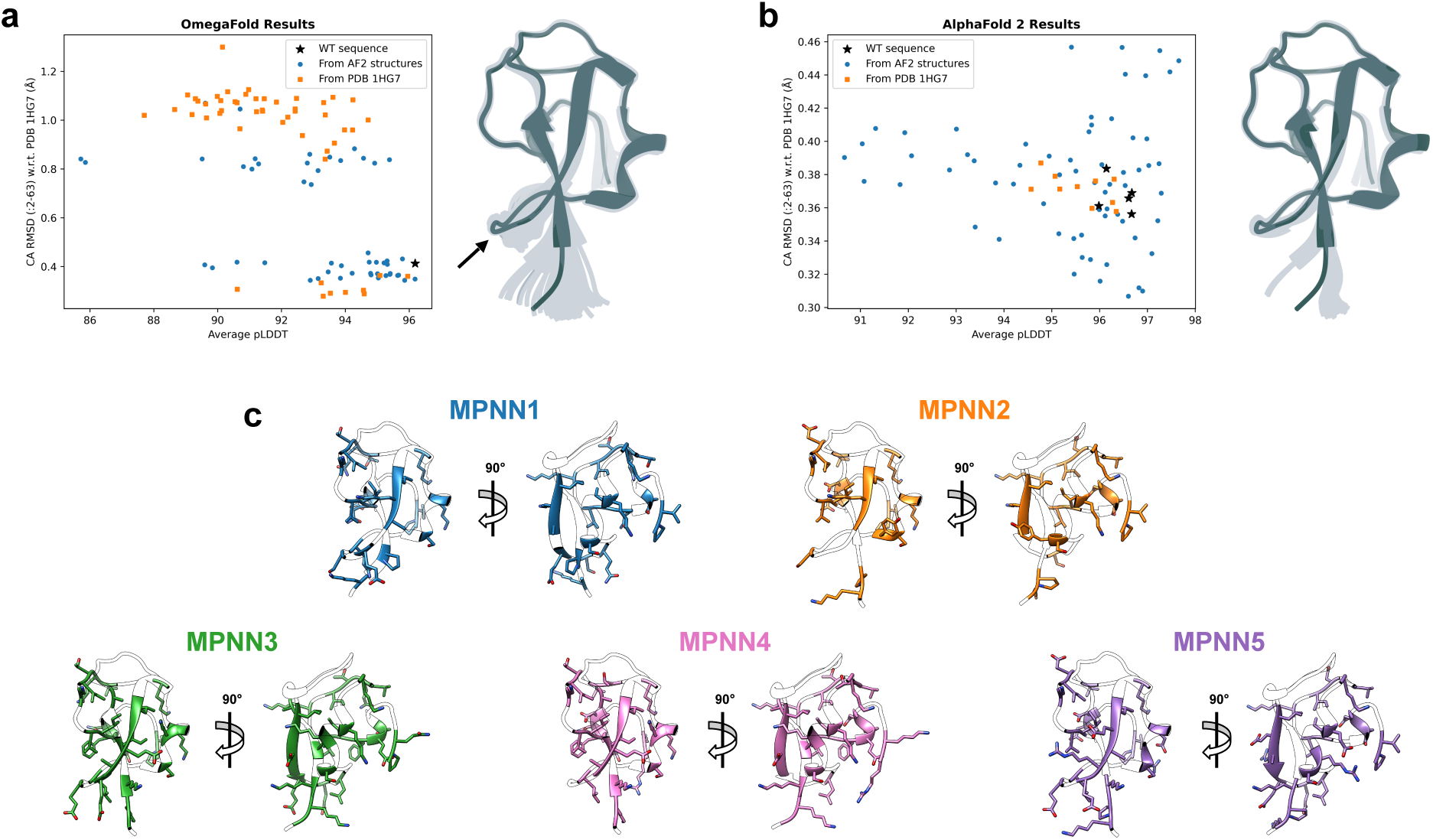
(a) OmegaFold results (alpha carbon RMSD versus average pLDDT) for the 100 designed sequences selected based on their global scores, with OmegaFold data for WT AFPIII included for comparison. To the right, all 100 OmegaFold structures of the Protein-MPNN sequences are shown in transparent gray overlaid on PDB 1HG7 (opaque); the arrow indicates the non-terminal loop with some structural variability. (b) AlphaFold 2 results (five structures per sequence) for the 15 sequences that had average pLDDT greater than 95 and RMSD less than 0.5 Å from OmegaFold, again with results for WT AFPIII included. To the right, the 75 AlphaFold structures of ProteinMPNN sequences are shown in transparent gray overlaid on PDB 1HG7 (opaque). (c) Two views of the AlphaFold rank 1 structure for each of the designs selected for expression. All residues that differ from the WT sequence are colored and have their side chains shown.

For the top 15 sequences, we next performed AF2 structure predictions as an additional screening step prior to selecting designs for expression. All 75 predicted AF2 structures (five models per sequence) closely resembled the crystal structure of AFPIII (PDB 1HG7) and had high confidence scores (Fig. 2b). We again applied the stringent cutoffs of average pLDDT greater than 95 and alpha carbon RMSD below 0.5 Å, calculated as the mean over all five AF2 structures for each sequence. Table 1 provides details about the ten sequences that passed this final filtering; the sequences are compared in Supplementary Fig. S1. These variants range from 58 to 70% sequence identity to WT AFPIII. Notably, all ten designs contain a higher number of charged residues than the WT. The variant labeled MPNN5 contains 15 charged residues, nearly double the eight present in WT AFPIII. In addition to their debated potential role in engulfment resistance, ^84^ we reasoned that charged residues may enhance the solubility and stability of the redesigned proteins.

**Table 1:** Top ten designs after screening with OmegaFold and AlphaFold 2.

| Label | Initial Structure | Sampling Temperature | % Identity<sup>a</sup> | # Charged Residues | Selected for Expression |
| --- | --- | --- | --- | --- | --- |
| MPNN1 | AF2 Rank 3 | 0.1 | 66.67 | 10 | Yes |
| MPNN2 | AF2 Rank 1 | 0.1 | 69.70 | 12 | Yes |
| MPNN3 | PDB 1HG7 | 0.2 | 57.58 | 12 | Yes |
| MPNN4 | PDB 1HG7 | 0.1 | 59.09 | 14 | Yes |
| MPNN5 | AF2 Rank 1 | 0.1 | 63.64 | 15 | Yes |
| MPNN6 | AF2 Rank 1 | 0.1 | 69.70 | 11 | No |
| MPNN7 | AF2 Rank 1 | 0.2 | 68.18 | 12 | No |
| MPNN8 | AF2 Rank 3 | 0.1 | 62.12 | 12 | No |
| MPNN9 | AF2 Rank 1 | 0.1 | 68.18 | 13 | No |
| MPNN10 | AF2 Rank 1 | 0.2 | 68.18 | 13 | No |

<sup>a</sup> With respect to rQAE m1.1 (PDB 1HG7).

We opted to express five of the top ten variants, aiming to balance experimental effort with the probability of obtaining informative designs. Because all ten candidates listed in Table 1 were predicted to adopt the correct fold with high confidence by both OF and AF2, structural confidence metrics were not used for further prioritization. Instead, we made a subjective selection based on the following factors: We chose the two sequences with the most charged residues (MPNN4 and MPNN5), the design with the fewest charged residues (MPNN1), the one with the lowest sequence identity to WT AFPIII (MPNN3), and the sequence with the lowest maximum percent identity (Supplementary Fig. S2) to any of the four already selected (MPNN2).

Fig. 2c shows the AF2 rank 1 models of the five selected designs, with substituted residues highlighted. Most of the altered residues are located on the protein’s surface, consistent with ProteinMPNN’s reported higher sequence recovery for buried residues.^9^ The mutations are widely distributed throughout the non-IBS regions rather than clustered in any specific location. Importantly, none of the modified side chains protrude substantially above the main IBS loop, which could have disrupted its flatness and potentially interfered with ice binding.

### Property predictions, expression outcomes, and CD spectra for AFPIII variants

Sumida et al. reported that redesigning myoglobin and the tobacco etch virus (TEV) protease with ProteinMPNN resulted in improvements in expression yield and thermostability, properties which can affect the practical utility of recombinant proteins for industrial applications.^17^ We expected that our AFPIII variants might likewise exhibit improvements in these traits. We therefore used computational tools to make predictions about these properties for the top ten sequences and compared those predictions to experimental outcomes for selected variants.

The tools NetSolP,^115^ SoluProt,^116^ and Protein-Sol^117^ were used to obtain predictions related to solubility and expression yield. These predictions were somewhat inconsistent between the different tools, with both NetSolP’s solubility scores and Protein-Sol’s scores higher for all ten designs than for WT AFPIII while SoluProt gave the highest score to the WT sequence (Fig. 3a). MPNN4 received the lowest score from SoluProt but the highest score from Protein-Sol. Meanwhile all ten designs and the WT received low usability scores from NetSolP.

**Figure 3:**
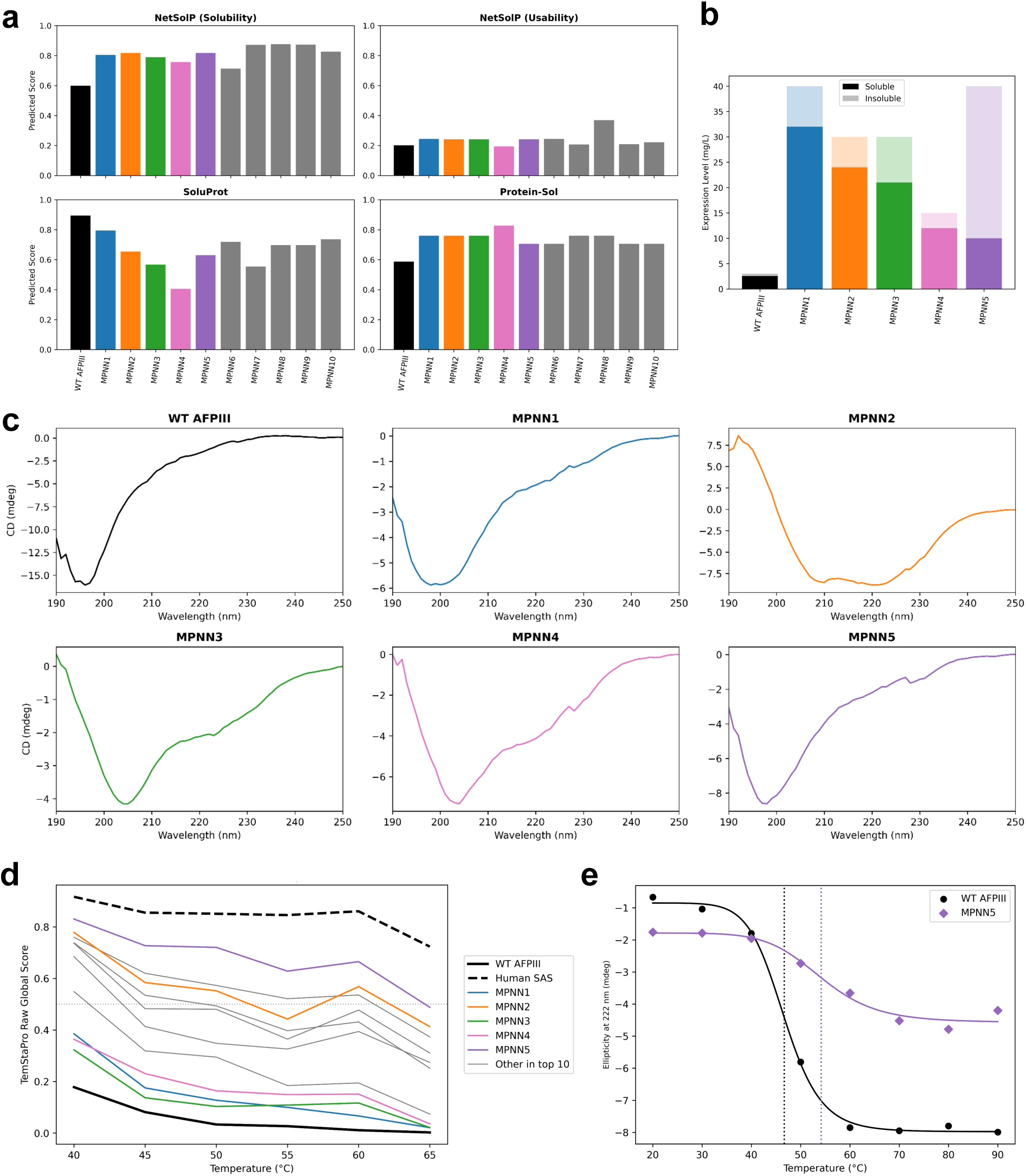
(a) NetSolP, SoluProt, and Protein-Sol scores for the top ten ProteinMPNN sequences, with WT AFPIII included for comparison. (b) Expression levels in *E. coli* after 16 hours at 15 °C for WT AFPIII and MPNN1-5 from GenScript’s expression evaluation, with the soluble fraction darker and the insoluble fraction lighter. (c) CD spectra at 20 °C for WT AFPIII and the five expressed variants. (d) Thermostability predictions (raw global score at each temperature) from TemStaPro for WT AFPIII, the top ten ProteinMPNN sequences, and the AFP-like domain of human sialic acid synthase. (e) Experimental thermostability data (ellipticity at 222 nm at different temperatures) for WT AFPIII and the MPNN5 variant, with solid lines representing sigmoidal c_1_u_0_rves fit to the data per Ref. 114 and dotted vertical lines at the EC50 values (46.7 °C and 54.2 °C, respectively).

To assess expressibility in *E. coli* for the five designs selected for experiments, we ordered a small-scale expression evaluation from GenScript, with WT AFPIII included for reference. Fig. 3b displays the expression levels from this evaluation, with the soluble fractions shaded darker. All five designed variants had higher soluble expression levels than the WT, with the largest difference for MPNN1 (40 mg/L and 80% soluble versus only 3 mg/L and 85% soluble for the WT). Separately from GenScript’s expression evaluation, we expressed and purified the selected variants MPNN1-5 for further characterization. Overall, the expression of the designed variants was high yielding, although the different yields among the variants did not clearly correlate with GenScript’s reported expression levels (Supplementary Fig. S5).

Fig. 3c shows the CD spectra of MPNN variants 1 to 5 and WT AFPIII at 20 *^◦^*C. WT AFPIII exhibited a pronounced minimum in molar ellipticity at ∼195 nm. Spectral deconvolution and fold recognition using the BeStSel web server ^118,119^ indicated a high antiparallel beta-sheet content with minimal helical contribution, consistent with the known crystal structure.^78^ The spectrum was also in good agreement with previously reported CD data for AFPIII.^109,120–125^

Among our five variants, the CD spectra suggest that two have structural features consistent with the WT fold, two others are somewhat different, and one is very different. MPNN1 and MPNN5 displayed CD spectra comparable to WT AFPIII, with minima in molar ellipticity between approximately 195 and 205 nm. Among these, MPNN5 most closely resembled the WT spectrum in both shape and magnitude. MPNN3 and MPNN4 were somewhat distinct from the WT in that the minimum in molar ellipticity was shifted to ∼205 nm. The structure of MPNN2 differed substantially from WT AFPIII and the other variants, as the spectrum showed an additional pronounced minimum near 222 nm. These spectral features indicate that MPNN2 has an increased helical contribution relative to WT AFPIII, whereas the other four variants consist primarily of beta-sheets with minimal helical content.

We used TemStaPro^126^ to predict the thermostability of the top ten sequences along with the WT. The AFP-like domain of human sialic acid synthase was included to provide a comparison to a mesophilic analog of the same fold.^127,128^ MPNN5 was predicted to have the greatest thermostability among the ProteinMPNN variants, closer to the scores for the human protein than to those for WT AFPIII (Fig. 3d). To experimentally assess thermostability, we monitored the ellipticity at 222 nm for MPNN5 and WT AFPIII while increasing the temperature from 20 *^◦^*C to 90 *^◦^*C in 10 degree increments. Fig. 3e shows that WT AFPIII exhibited a melting temperature of ∼47 *^◦^*C, in line with previous studies.^121–123^ We found that MPNN5 displayed a higher apparent melting temperature of ∼54 *^◦^*C, indicating moderately enhanced thermostability relative to WT AFPIII. TemStaPro was thus correct in its ranking of the two proteins’ stability, but the magnitude of the enhancement was smaller than the predictions suggested.

### Thermal hysteresis activity and ice crystal morphology

Next, we evaluated the thermal hysteresis activity of the designed variants at approximately 50 *µ*M and compared them to WT AFPIII using a Nanoliter Cryoscope (Fig. 4). In solutions containing WT AFPIII, a ∼10 *µ*m disc-shaped seed ice crystal transitions into a blunt hexagonal bipyramid when held slightly below its melting temperature (Fig. 4b). With incremental cooling, this morphology sharpens into a well-defined hexagonal bipyramid characterized by a *c*-to-*a* ratio of ∼1.5. No further growth is apparent until reaching a point where the bound AFPIII can no longer constrain the ice crystal’s growth and typically a single, fine spicule propagates from the tips of the hexagonal bipyramid, which appear to thicken slightly. Several additional spicules then grow rapidly, parallel to the initial ones; this burst of spicular growth is described as the nonequilibrium freezing point. The gap between this burst temperature and the melting temperature is referred to as thermal hysteresis.

**Figure 4:**
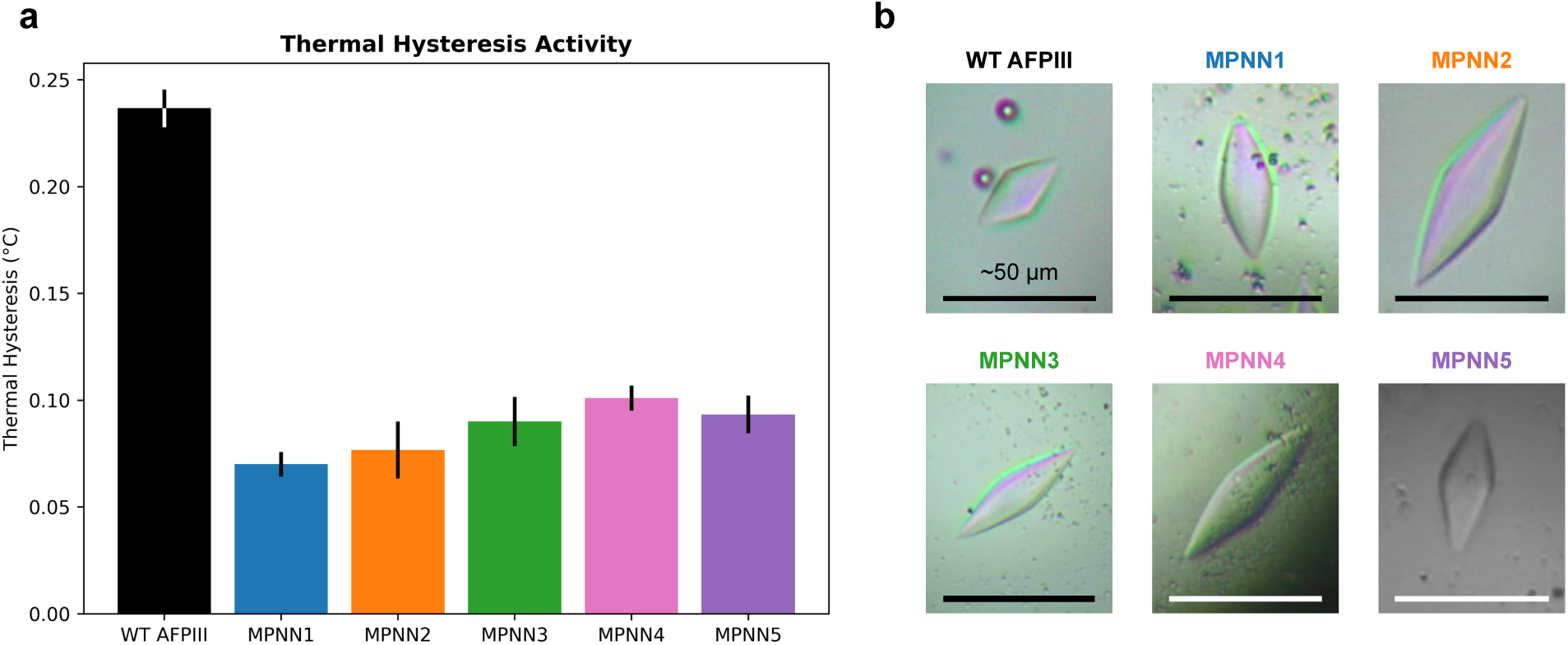
(a) Thermal hysteresis activity of WT AFPIII and the ProteinMPNN variants, with error bars indicating the standard error from three replicates. (b) Ice crystals in the presence of WT AFPIII and the variants, in a buffer of 50 mM Tris/150 mM NaCl (pH 7.5). Ice crystals were photographed at the time of tip elongation.

All five ProteinMPNN designs exhibited measurable thermal hysteresis, with their activity around 30 to 40% of that observed for WT AFPIII (Fig. 4a). Initial crystal shaping near the melting point was similar to the WT protein’s effect, with disc-shaped seeds converting to blunt and then regular hexagonal bipyramids. Continued cooling did not result in significant growth arrest. Instead, crystals elongated progressively along the *c*-axis, with the tips extending toward the liquid interface while maintaining nearly constant lateral dimensions. Once the advancing tip reached the interface, further elongation along the *c*-axis ceased and growth proceeded along the *a*-axes, yielding a single thick spear-like crystal without additional temperature reduction. The resulting growth behavior resembles that observed for smaller antifreeze glycoprotein (AFGP) isoforms and other AFPs with relatively weak ice-binding activity.^129^ Meanwhile screening with a gold nanoparticle (AuNP) colorimetric assay^130^ indicated that our designs vary in their effects on AuNP aggregation during a freeze-thaw cycle, potentially reflecting altered ice recrystallization inhibition (IRI) activity relative to WT AFPIII (Supplementary Fig. S6).

## Conclusions

Recent advances in deep learning-based protein design present new possibilities for redesigning natural AFPs. Our fixed-backbone computational redesign of AFPIII produced variants that expressed efficiently in soluble form. With minimal customization in the design process, we obtained AFPIII variant sequences confidently predicted to adopt the correct fold, and expressing five variants was sufficient to produce a subset displaying secondary structure signatures consistent with the WT protein. These observations suggest that AFPIII’s structure is relatively designable with ProteinMPNN.

Our designed AFPIII variants exhibited properties desirable for practical applications, namely high soluble expression levels and, in one case, improved thermostability relative to WT AFPIII. Interestingly, despite preserving all residues previously identified as part of the canonical ice-binding site,^59,80^ even the variants that appeared to maintain AFPIII’s structural features showed reduced thermal hysteresis activity. It is possible that our designs possessed weakened affinity for ice due either to the loss of previously unrecognized residues that are involved in the WT protein’s binding or to the addition of disruptive side chains. Alternatively, we speculate that residues outside the established ice-binding surface may contribute to antifreeze activity by modulating the structure and dynamics of hydration water, conferring an optimal amount of backbone rigidity, or resisting engulfment.

While our present efforts did not produce variants that simultaneously maintained full activity while improving other traits such as thermostability, this goal may be well within the reach of current protein design methods. Either rationally selecting individual residues in our current variants to revert to the WT sequence or instead generating new sequences with more positions fixed may provide an avenue to preserve the full activity. We anticipate that our original set of fixed residues can be expanded by incorporating additional sites based on proximity to the reported IBS or conservation among related natural AFPs.

The toolkit for computational protein design has expanded dramatically in recent years, with these deep learning innovations only beginning to permeate the field of ice-binding proteins.^111,112^ Our AFPIII designs emphasize the value of cutting-edge computational methods for producing modified AFPs/IBPs with properties distinct from those found in nature. In addition, our results underscore the need for further exploration of the functional impact of modifications outside of AFPs’ reported ice-binding regions, a consideration which will likely influence the outcomes of future AFP (re)design and optimization efforts.

## Methods

### ProteinMPNN

For each of the six starting structures (PDB 1HG7^78^ and five AlphaFold^1,113^ structures of rQAE m1.1^58^), we ran the v_48_020 ProteinMPNN model^9^ with sampling temperatures 0.1, 0.2, and 0.3 (50 sequences per temperature per structure). Cysteines were prohibited in the AFPIII variant designs. In addition to the N-terminal methionine, the following residues were kept fixed based on the ice-binding surfaces described in Ref. 80: Gln9, Leu10, Pro12, Ile13, Asn14, Thr15, Ala16, Thr18, Leu19, Val20, Met21, Val41, Ser42, and Gln44. We also carried out an in silico test applying a similar redesign approach to a different fish AFP; the methods and results of this test are included in the Supplementary Information. We produced sequence logos using WebLogo.^131^

### Structure prediction and analysis

We ran OmegaFold^4^ with ten cycles. AlphaFold predictions were made using ColabFold v1.5.5^113^ with three recycles, default multiple sequence alignment mode (MMseqs2 UniRef+ Environmental^132,133^), no templates, no relaxation, and all five AlphaFold 2 ptm models. A test using AlphaFold in single-sequence mode had shown that running without multiple sequence alignments produced low-quality predictions for WT AFPIII and all 100 designed sequences (Supplementary Fig. S3). We assessed predicted structures with Biopython. ^134^ Average pLDDT was calculated over all 66 residues, whereas the alpha carbon RMSD with respect to PDB 1HG7 was calculated excluding two residues at each terminus. Structures were visualized using UCSF Chimera.^135^

### Property predictions

Predictions pertaining to expression and solubility were made using the web servers of NetSolP (https://services.healthtech.dtu.dk/services/NetSolP-1.0/),^115^ SoluProt (https://loschmidt.chemi.muni.cz/soluprot/),^116^ and Protein-Sol (https://protein-sol.manchester.ac.uk/).^117^ The NetSolP predictions included both solubility and usability, and the model type was ESM1b.^136^ Thermostability predictions were made with TemStaPro^126^ using Neurosnap’s online platform.^137^ The AFP-like segment of human sialic acid synthase included for comparison was KLGKSVVAKVKIPEGTILTMDMLTVKVGEPKGYPPEDIFNLVGKKVLVTVEEDDTIMEELVDNHGK.^127^

### Expression evaluation in *E. coli*

We employed GenScript’s expression evaluation service to assess the expressibility of WT AFPIII and the five designs, each with a C-terminal hexahistidine (His6) tag appended. All expressed sequences are listed in Supplementary Table S1. The template vector was pET30a and the restriction sites were NdeI (CATATG) and HindIII (AAGCTT). GenScript transformed *Escherichia coli* (*E. coli*) BL21 Star™ (DE3) competent cells with the plasmid, inoculated LB medium containing kanamycin, and incubated the cultures at 37 °C and 200 rpm. The expression volume was 4 mL. Expression was induced with 0.5 mM IPTG when the cell density reached OD 0.6-0.8 at 600 nm. GenScript reported that expression for 16 hours at 15 °C appeared more favorable than four hours at 37 °C for all six proteins. SDS-PAGE gels are shown in Supplementary Fig. S4.

### Expression and purification of designed variants

Separately from GenScript’s expression evaluation, we expressed and purified the five designed variants to obtain samples for structure and activity characterization. The genes coding for the MPNN1-5 variants, with an added C-terminal His6 tag, were synthesized by Twist Bioscience and cloned into a pET28 expression vector, using the Xho1/Xba1 restriction sites. The plasmid containing the corresponding gene was transformed into competent *E. coli* BL21 (DE3) cells and plated on LB-agar plates supplemented with kanamycin. A single colony was selected and used to inoculate 100 mL starter culture of LB in the presence of 50 *µ*g/mL kanamycin. The culture was grown overnight under continuous shaking at 37 °C. The following day the dense starter culture was used to inoculate (1.5%) flasks with 1 L LB, in the presence of kanamycin. The cultures were grown at 37 °C under continuous shaking until OD_600_ reached 0.6-0.8, after which the temperature was reduced to 16 °C. Subsequently, 0.5 mM IPTG was added to initiate the overnight expression of the target protein. Cells were collected by centrifugation at 2000 rpm for 30 minutes and the cell pellet was stored at -20 °C until further processing.

The C-terminal His6 tag enabled the purification of the target protein by immobilized metal affinity chromatography (IMAC). The cell pellet with the target protein was thawed on ice, resuspended in lysis buffer (50 mM Tris, 150 mM NaCl, pH 7.5) and sonicated to disrupt the cells. The cell lysate was centrifuged for 1 hour at 4700 rpm, separating the cell debris from the clear lysate. Ni-NTA resin (HisPur™, Thermo Fisher) was applied to a gravity column and washed with water, followed by equilibration with the lysis buffer. The supernatant of the centrifuged cell lysate was applied to the equilibrated Ni-NTA column and the flow-through was collected. The Ni-NTA resin was subsequently washed with 10 column volumes of lysis buffer. The target protein was eluted by a step increased gradient of imidazole supplemented lysis buffer, with final concentrations of imidazole ranging between 5 and 250 mM and each with a volume of 4 mL. The collected fractions were checked by SDS-PAGE (15%) (Supplementary Fig. S5) and fractions containing pure target protein were pooled and dialyzed overnight in 3.5k MWCO dialysis tubing (SnakeSkin™, Thermo Fisher) against lysis buffer. Subsequently, the dialyzed protein solution was concentrated with a 3k centrifugal concentrator (Amicon^®^, MilliporeSigma) at 3900 rpm. The concentration of the target protein was determined by a Bradford protein assay (Pierce™, Thermo Fisher) against a BSA standard. Additional details about the materials used in this process are included in the Supplementary Information.

### Circular dichroism

Samples were prepared for circular dichroism by dialysis (3.5K MWCO Snakeskin Dialysis Tubing, 16 mm dry I.D., 35 feet, Thermo Scientific, IL, USA) against pure water for 48 hours with three changes of water. 200 *µ*L of dialyzed sample was loaded into a cuvette with a pathlength of 1 mm. A Jasco J-1500 Spectropolarimeter (JASCO Inc., MD, USA) equipped with a temperature control unit interfaced to a computer was used to obtain protein secondary structures. Each scan was performed continuously from 180-250 nm with a scanning speed of 100 nm/min. The data pitch was 0.1 nm and the accumulation was set to 3. Pure water and ultrapure WT AFPIII (QAE with the His-tag removed^70^) served as the negative and positive controls, respectively. Thermostability measurements of MPNN5 and AFPIII were performed from 20 °C to 90 °C in 10 degree increments. The BeStSel webserver^118,119^ was used for secondary structure determination and fold recognition. The wavelength range was defined as 190-250 nm with a scale factor of 1.

### Thermal hysteresis

Thermal hysteresis, defined as the temperature difference (°C) between the melting and freezing points of a solution, was measured using a Clifton Nanoliter Osmometer. A specialized cold stage equipped with an internal water circulation as a heat sink and a separate pipeline for dry air was used. A droplet of immersion oil B (Cargille Laboratories, Cedar Grove, NJ, freezing point *<* -13 °C) was placed on the underside of a 6-cell copper sample holder. ^138,139^ The sample holder was positioned in the cold stage under an optical microscope with 100x magnification (Zeiss Axioscope 5, Carl Zeiss Microscopy GmbH, Jena, Germany). A buffer of 50 mM Tris and 150 mM NaCl (pH 7.5) was prepared. The protein samples were brought to the following concentrations in buffer: MPNN1 (23.4 *µ*M), MPNN2 (50 *µ*M), MPNN3 (50 *µ*M), MPNN4 (50 *µ*M), and MPNN5 (45 *µ*M). An ultrapure solution of WT AFPIII^70^ (50 *µ*M) was used as a positive control. Glass capillaries were prepared using a pipette puller. Samples were loaded into the tip of the capillary and carefully injected into the center of one of the holes in the sample holder. Prior to cooling, the sample holder was covered with a glass cover slip to prevent moisture from accessing the sample. To obtain ice crystals, the whole sample was frozen and then melted back until one crystal remained. Temperatures were manually adjusted using the control knob on the Clifton Nanoliter Osmometer. Thermal hysteresis measurements were performed with a cooling rate of 0.075 °C/min and with a two minute annealing time for each sample. Measurements were performed three times per sample. The copper disk was thoroughly cleaned with chloroform and water before and after each use.

### Gold nanoparticle colorimetric assay

Spherical gold nanoparticles (AuNPs) with a diameter of 10 nm and functionalized with nitrilotriacetic acid modified thioctic acid (Au10-TA-NTA) were synthesized as previously reported.^130,140,141^ We employed the AuNP colorimetric assay and analyzed the data as previously described.^130^

## Acknowledgement

C.N.C. is grateful to Keaun Amani for providing credits on the Neurosnap platform. A.J.A. thanks Dr. Graham Heberlig for technical support and advice on cloning strategies. We also thank Juliana Tang, Ruilin Zhao, and Crystal Li for their assistance with GenScript’s expression services. This work was supported by the Air Force Office of Scientific Research under awards nos. FA9550-20-1-0351 and FA9550261B096. C.N.C. was partially supported by a San Diego Fellowship.

## Supporting Information Available

The Supporting Information is available free of charge and includes the following details: materials used for expressing the variants, the results of the AuNP colorimetric assay screening, the methods and results from a redesign test using a different fish AFP, additional information comparing the top ten designed AFPIII sequences, AlphaFold 2 single sequence mode prediction outcomes, SDS-PAGE gels, and all expressed sequences.

## Data availability

All data will be made publicly available at the time of publication.

## Code availability

The main tools used in this study, including ProteinMPNN, ColabFold, and OmegaFold, are publicly available. Analysis scripts in Python are available from the authors upon request.

## TOC Graphic

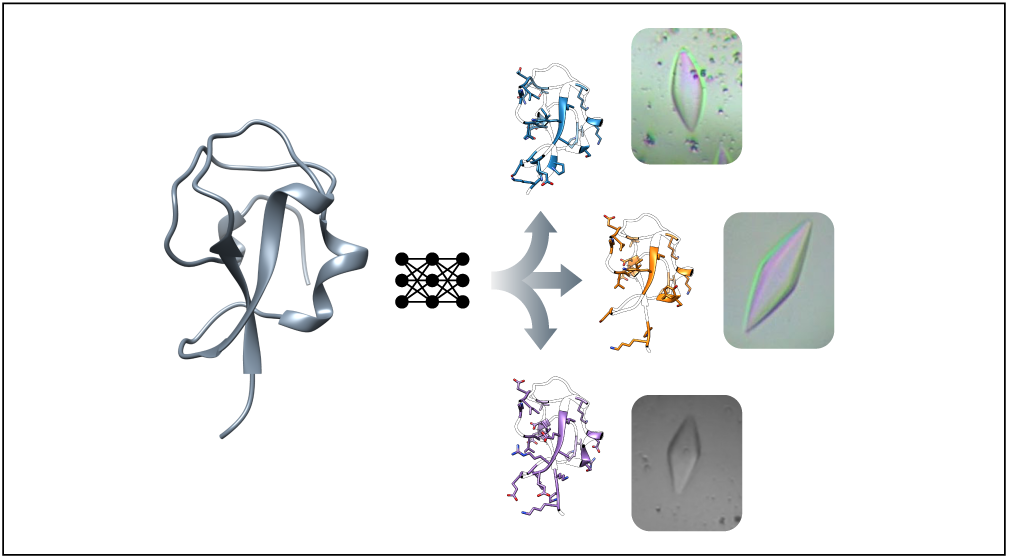

