## Supporting Information for "Computational Redesign of an Antifreeze Protein Using Deep Learning"

### Supporting Information for:
### Computational Redesign of an Antifreeze Protein Using Deep Learning

Cianna Calia,[1,a)] Arthur J. Altunc,[1] Rosemary J. Eufemio,[2] Benjamin O. Alvarado,[1] Jake D. Huynh,[1] Eunkeu Oh,[3] Michael D. Burkart,[1] Konrad Meister,[2,b)] and Francesco Paesani[1,4,5,6,c)]

[1)] *Department of Chemistry, University of California San Diego, La Jolla, California 92093, United States*

[2)] *Department of Chemistry and Biochemistry, Boise State University, Boise, Idaho 83725, United States*

[3)] *Optical Sciences Division, U.S. Naval Research Laboratory, Washington, District of Columbia 20375, United States*

[4)] *Materials Science and Engineering, University of California San Diego, La Jolla, California 92093, United States*

[5)] *Halicioğlu Data Science Institute, University of California San Diego, La Jolla, California 92093, United States*

[6)] *San Diego Supercomputer Center, University of California San Diego, La Jolla, California 92093, United States*

#### CONTENTS

I. **Materials for variant expression and purification** S2

II. **AuNP colorimetric assay** S2

III. **In silico redesign test using a sea raven AFP structure** S2

[a)] Electronic mail:

[b)] Electronic mail:

[c)] Electronic mail:

#### I. MATERIALS FOR VARIANT EXPRESSION AND PURIFICATION

Chemicals and reagents used for expression and purification were purchased from the following commercial sources: Miller's Luria Broth (LB, granulated) and agar (molecular biology grade, high gel strength) were purchased from Research Products International (RPI). Kanamycin sulfate, isopropyl $\beta$-D-1-thiogalactopyranoside (IPTG) and Tris base were purchased from Fisher Bioreagents. Competent *E. coli* cells, DH5$\alpha$ and BL21 (DE3), were purchased from New England Biosciences (NEB). Sodium chloride (NaCl) and sodium dodecyl sulfate (SDS) were purchased from Fisher Scientific. Ni-NTA resin (HisPur™) and unstained protein molecular weight marker (Pierce™) were purchased from Thermo Fisher. Phosphate-buffered saline (PBS 10x, OmniPur®) was purchased from EMD Millipore. Ultrapure water (Sartorius Arium® Pro) was used to dissolve chemicals and reagents. Buffers were filtered using a bottle-top vacuum filtration system (Nalgene® DS0320-2545, Thermo Fisher) equipped with a 0.2 $\mu$m mixed cellulose ester membrane filter (Whatman, Cytiva).

Equipment and consumables were employed from the following sources: A Kühner incubator shaker (ISF-1-W) was used for bacterial growth of *E. coli*. A Beckman Coulter Avanti™ J-20 XP centrifuge was used to collect *E. coli* cells through centrifugation with a JLA-8.1 rotor. An Ultrasonics Sonic processor FS-600N was used to sonicate resuspended cell pellets. SnakeSkin™ dialysis tubing MWCO 3.5k was purchased from Thermo Scientific and used for dialysis of the purified target protein. Centrifugal concentrators (3k, Amicon®) were purchased from MilliporeSigma. An Eppendorf Centrifuge 5804 R (15 amp version) and a Thermo Scientific Heraeus Multifuge X3R were used to spin concentrate purified target protein.

#### II. AUNP COLORIMETRIC ASSAY

The gold nanoparticle (AuNP) colorimetric assay employs functionalized AuNPs as an indirect probe to screen for potential ice recrystallization inhibition (IRI) activity in a microwell plate format. The AuNP dispersion, which is red of color, is titrated along a two-fold dilution series of the target protein and subjected to a freeze-thaw cycle. In the absence of any IRI-active agents, the whole mixture is frozen, forcing the AuNPs to aggregate. This aggregation results in a color change to blue, due to the surface plasmon resonance (SPR) of the AuNPs. The color change is by design irreversible, even when the microwell plate is thawed to room temperature. In the presence of an IRI agent the recrystallization of the ice crystals is inhibited, helping the AuNPs to stay in their monodisperse state and maintain their red color[1–3]. The change in color is easily detected by measuring the absorbance spectra before and after the freeze-thaw cycle; subsequent calculation of the "change in monodispersity" and pharmacology-inspired analysis allows quantification through $IC_{50}$ values[3].

We analyzed our ProteinMPNN variants with this assay as recently described[3] (Supplementary Fig. S6a). Only MPNN4 and to a lesser degree MPNN3 produced relatively well-defined sigmoidal curves (Supplementary Fig. S6b). These variable outcomes, distinct from the reported curve for WT AFPIII[3], suggest that the sequence modifications in some of our variants affected their ability to modulate the behavior of AuNPs during a freeze-thaw cycle and may potentially reflect changes in IRI activity.

#### III. IN SILICO REDESIGN TEST USING A SEA RAVEN AFP STRUCTURE

As an initial in silico test of the generalizability of the fixed-backbone redesign approach to another non-repetitive AFP structure, we also produced ProteinMPNN variants of the sea raven type II AFP (Supplementary Fig. S7). We used a temperature of 0.1 to generate 150 sequences based on model 1 from the NMR structure PDB 2AFP[4]. Cysteines were excluded in 50 of the sequences, allowed in another 50, and both fixed and allowed in the remaining 50. In all cases we fixed residues Thr91, Lys92, Pro93, Asp94, Asp95, Val96, Leu97, Ala98, Ala99, and Ser120 based on their reported importance for this AFP's thermal hysteresis activity[5].

Out of the 150 sequences we generated, none achieved an average pLDDT value greater than 80 from either OF or AF2, and the WT sequence had disparate average pLDDT values from the two models. This preliminary test suggests that redesigning the type II AFP structure may present distinct challenges that, at a minimum, require further exploration of possible ProteinMPNN settings, starting structures, and structure prediction models. We did not express any of these sequences.

```
    WT: MNQASVVANQLIPINTALTLVMMRSEVVTPVGIPAEDIPRLVSMQVNRAVPLGTTLMPDMVKGYAA
 MPNN1: *K*P******************LKA*E*S*Q****S***K***K**TQTI*K**VIQ*E*****VP
 MPNN2: *KP*******************LKG***S*****YS***K***K**V*DI*K**VI**ED****VP
 MPNN3: *GKK*I**A*************LGEAE*E*L****S*K*K***K**VQTI*K**VIFEN*****QP
 MPNN4: *GKK*I**A*************LGEA****L****S*K*K***K*TV*TI*K**VIFEND****KR
 MPNN5: *KPK******************LEARE*E*L****SR**E***K**V*DI*K**VI**ED****VP
 MPNN6: *KP*******************LKA***E******SR**E***K**VQDI*K**VI**E*****VP
 MPNN7: *KP*******************LEAR*********SR**E***K**V*DI*K**VI**ED***FVP
 MPNN8: *KEP*I****************LKA*Q*Y*Q****S***K***K**T*TI*K**VIQAED****VP
 MPNN9: *KP*******************LKA*Q*E******SR**E***K**V*DI*K**VI**ED****VP
MPNN10: *KEP******************LKA***S*L****S***K***K**V*DI*K**VI**ED****VP
```

FIG. S1. The top ten ProteinMPNN sequences after screening with OmegaFold and AlphaFold 2, compared to WT AFPIII. The WT sequence refers to rQAE m1.1. Asterisks indicate residues in each variant that match the WT sequence. Fixed residue positions are shaded in gray.

| | WT | MPNN1 | MPNN2 | MPNN3 | MPNN4 | MPNN5 | MPNN6 | MPNN7 | MPNN8 | MPNN9 | MPNN10 |
|---|---|---|---|---|---|---|---|---|---|---|---|
| WT: | 100.00 | 66.67 | 69.70 | 57.58 | 59.09 | 63.64 | 69.70 | 68.18 | 62.12 | 68.18 | 68.18 |
| MPNN1: | 66.67 | 100.00 | 83.33 | 75.76 | 68.18 | 80.30 | 84.85 | 77.27 | 89.39 | 81.82 | 87.88 |
| MPNN2: | 69.70 | 83.33 | 100.00 | 69.70 | 71.21 | 84.85 | 89.39 | 87.88 | 81.82 | 90.91 | 92.42 |
| MPNN3: | 57.58 | 75.76 | 69.70 | 100.00 | 89.39 | 74.24 | 72.73 | 66.67 | 72.73 | 69.70 | 72.73 |
| MPNN4: | 59.09 | 68.18 | 71.21 | 89.39 | 100.00 | 71.21 | 66.67 | 69.70 | 72.73 | 68.18 | 74.24 |
| MPNN5: | 63.64 | 80.30 | 84.85 | 74.24 | 71.21 | 100.00 | 89.39 | 92.42 | 78.79 | 92.42 | 87.88 |
| MPNN6: | 69.70 | 84.85 | 89.39 | 72.73 | 66.67 | 89.39 | 100.00 | 90.91 | 78.79 | 95.45 | 87.88 |
| MPNN7: | 68.18 | 77.27 | 87.88 | 66.67 | 69.70 | 92.42 | 90.91 | 100.00 | 77.27 | 92.42 | 86.36 |
| MPNN8: | 62.12 | 89.39 | 81.82 | 72.73 | 72.73 | 78.79 | 78.79 | 77.27 | 100.00 | 83.33 | 87.88 |
| MPNN9: | 68.18 | 81.82 | 90.91 | 69.70 | 68.18 | 92.42 | 95.45 | 92.42 | 83.33 | 100.00 | 89.39 |
| MPNN10: | 68.18 | 87.88 | 92.42 | 72.73 | 74.24 | 87.88 | 87.88 | 86.36 | 87.88 | 89.39 | 100.00 |

FIG. S2. Percent identity matrix for WT AFPIII (rQAE m1.1) and the top ten ProteinMPNN sequences.

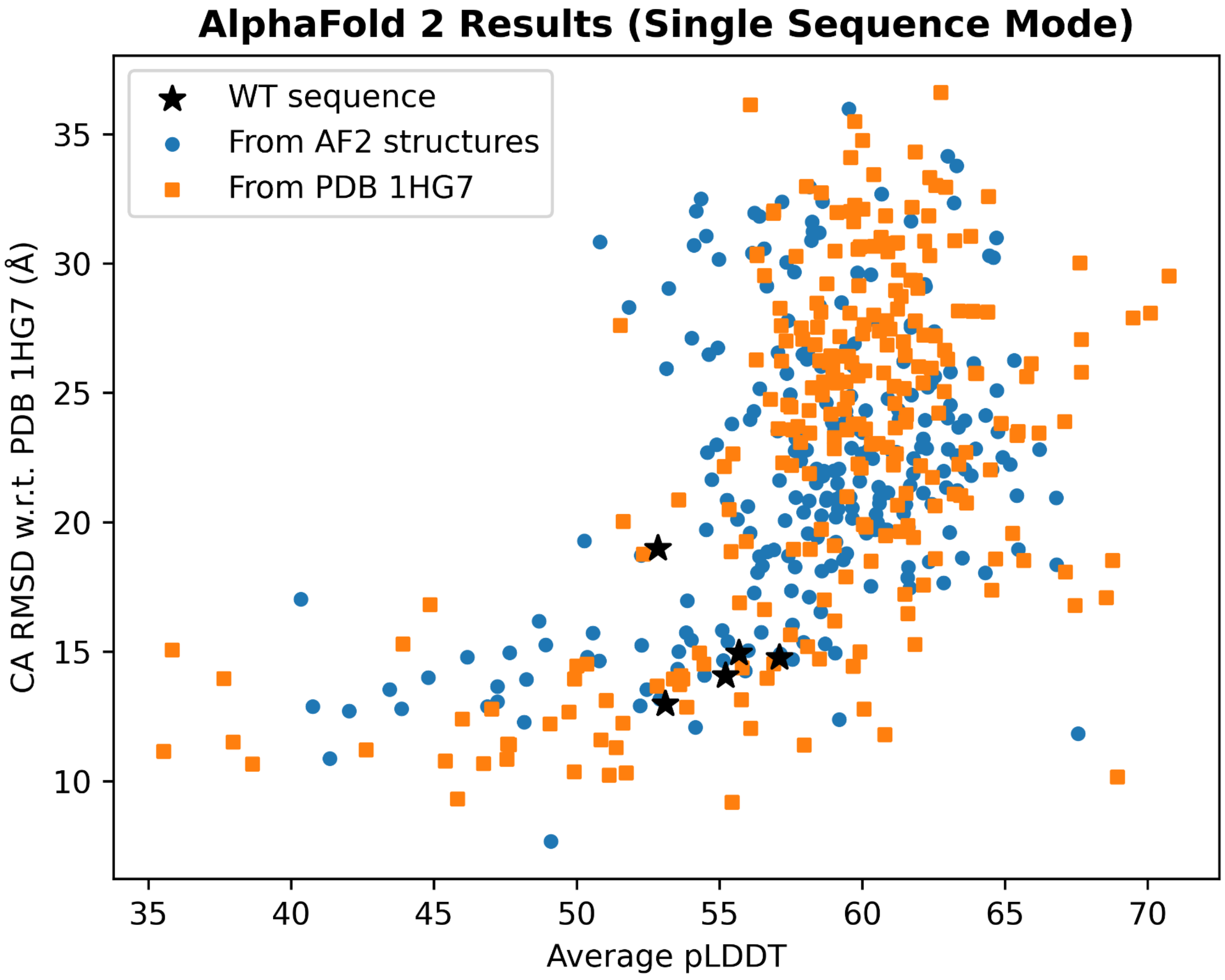


FIG. S3. AlphaFold 2 single sequence mode results (alpha carbon RMSD versus average pLDDT for all residues) for the 100 designed sequences selected based on their global scores, with analogous results for WT AFPIII included for comparison.

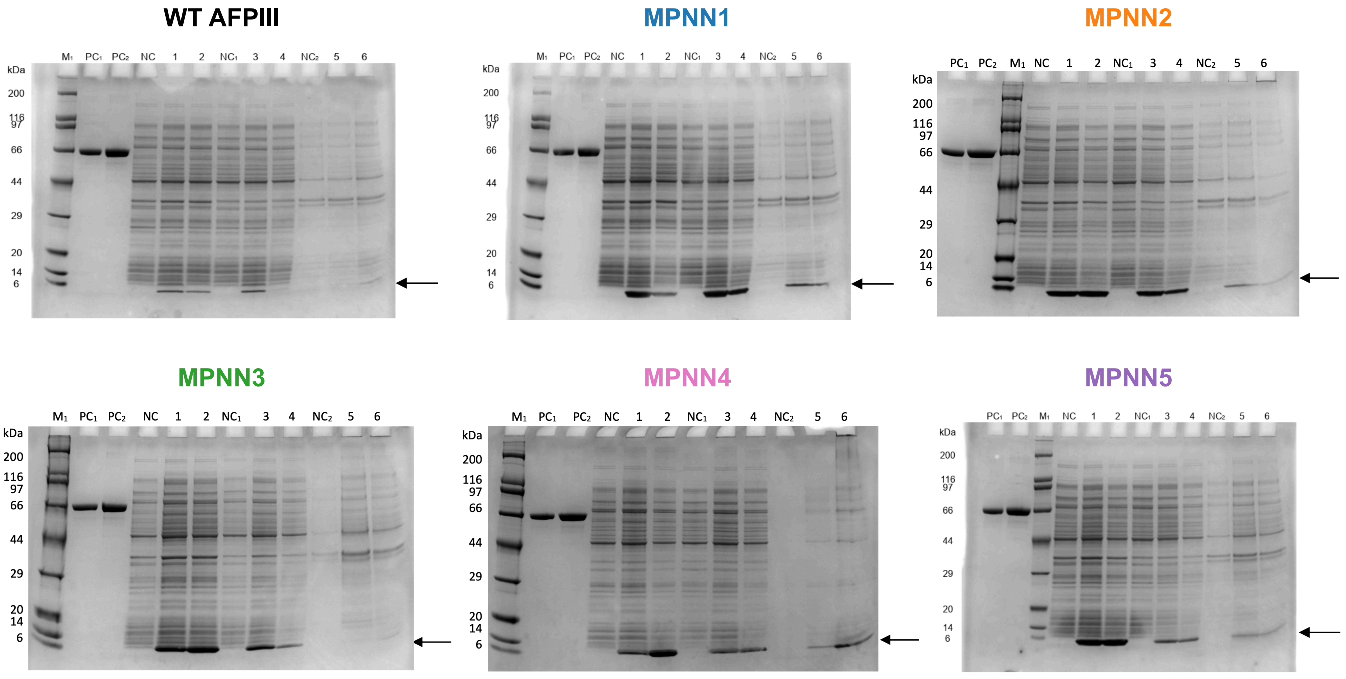


FIG. S4. SDS-PAGE gels from GenScript's expression evaluation for WT AFPIII and the five designed variants. The lanes are labeled as follows: $M_1$: protein marker, $PC_1$: BSA (1 $\mu$g), $PC_2$: BSA (2 $\mu$g), NC: cell lysate without induction, 1: cell lysate with induction for 16 hours at 15 °C, 2: cell lysate with induction for 4 hours at 37 °C, $NC_1$: supernatant of cell lysate without induction, 3: supernatant of cell lysate with induction for 16 hours at 15 °C, 4: supernatant of cell lysate with induction for 4 hours at 37 °C, $NC_2$: pellet of cell lysate without induction, 5: pellet of cell lysate with induction for 16 hours at 15 °C, and 6: pellet of cell lysate with induction for 4 hours at 37 °C.

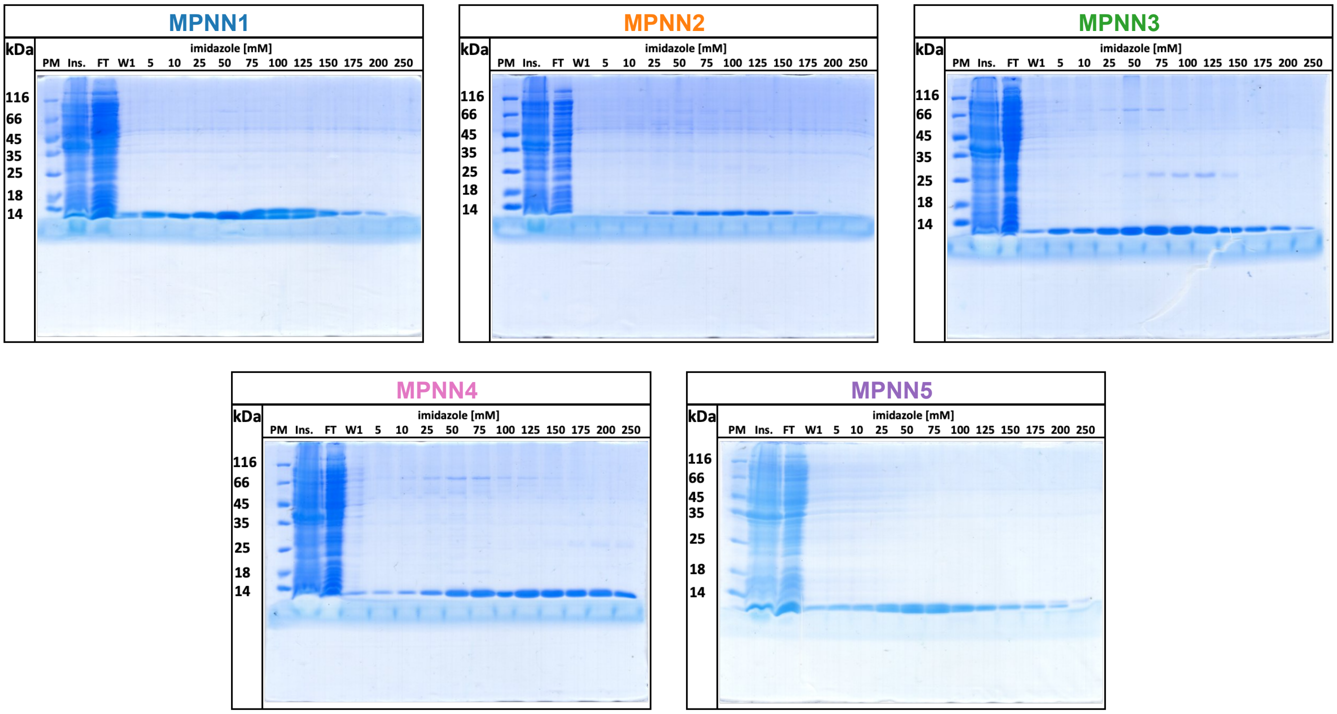


FIG. S5. SDS-PAGE (15%) gels after IMAC (Ni-NTA) purification of the five variants we expressed. Shown from left to right: protein marker (PM) in kDa, insoluble fraction (Ins.; pellet after sonication and centrifugation), and flow through (FT) of the supernatant. Wash 1 (W1) is the elution of the protein with lysis buffer, followed by increasing concentration of imidazole starting from 5 mM up to 250 mM.

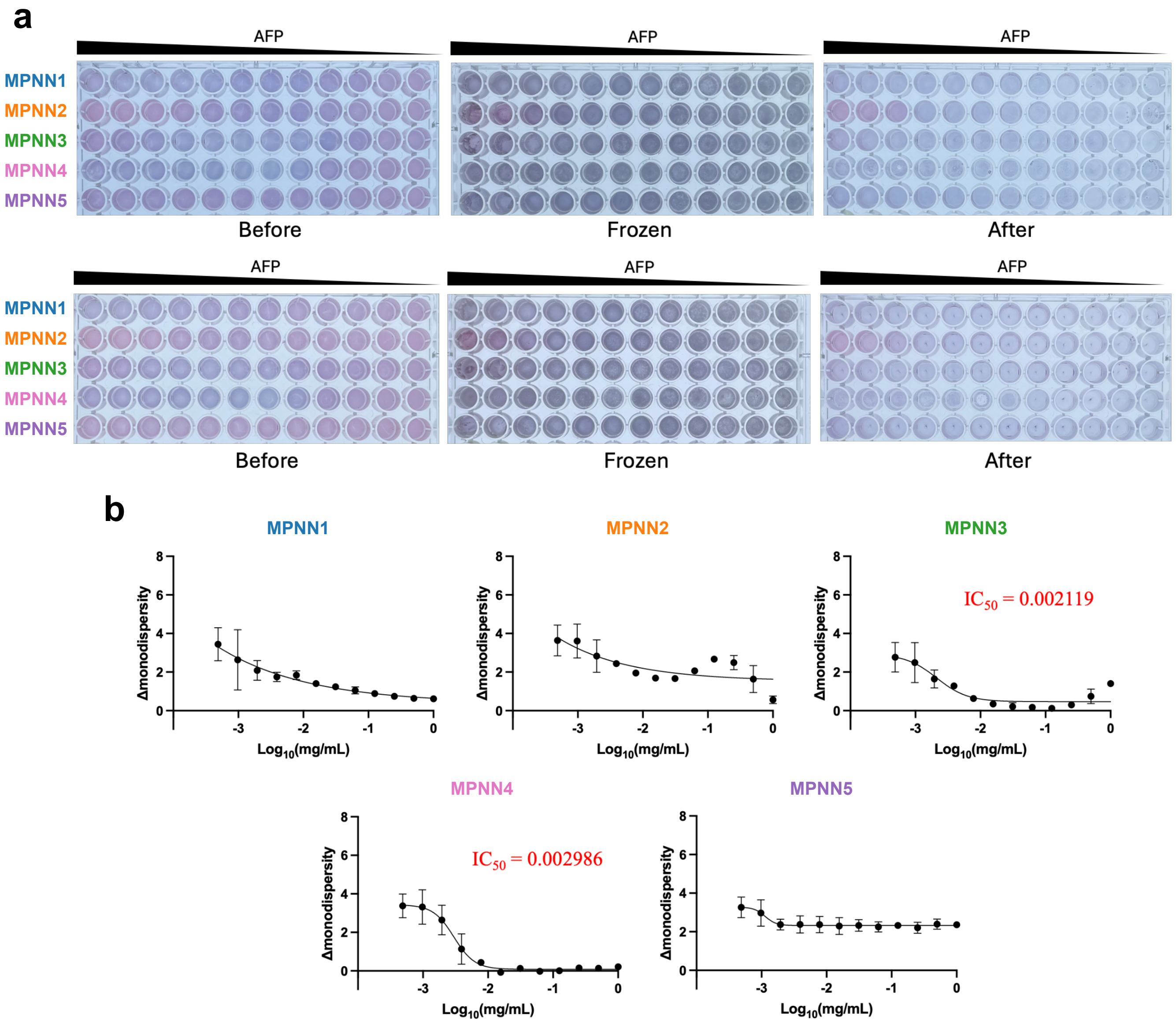


FIG. S6. (a) AuNP colorimetric assay 96-microwell plate layout at different timepoints of the freeze-thaw cycle for duplicate screenings (Screening 1 on top and Screening 2 below). MPNN variants were subjected to a two-fold serial dilution, starting at 1 mg/mL. "Before" refers to the state upon mixing the protein with the AuNPs. "Frozen" refers to the frozen state after 2 hours of freezing. "After" refers to the state after thawing. (b) Hill plots from the AuNP colorimetric assay for the five variants. The change in monodispersity is expressed as a function of the concentration in mg/mL on a logarithmic scale. $IC_{50}$ values in mg/mL are shown in red for the two variants with relatively well-defined sigmoidal trends. Solid lines represent the Hill plot fitting. Each protein was tested in duplicate, and the average values are shown. An analogous plot and an $IC_{50}$ value of 0.103 mg/mL were reported previously for WT AFPIII[3].

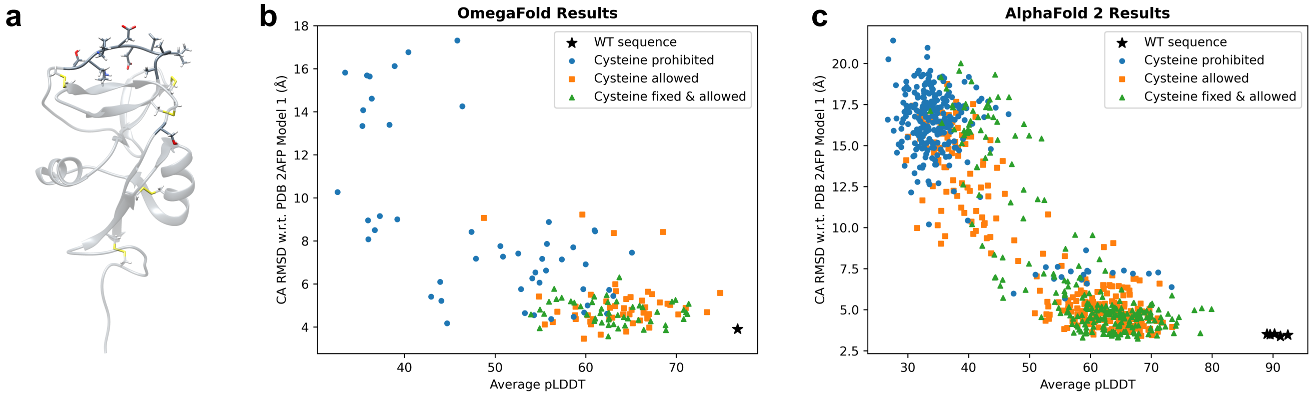


FIG. S7. (a) Structure of the sea raven type II AFP that was provided to ProteinMPNN (PDB 2AFP model 1[4]), with side chains shown for residues that were fixed based on reported functional significance[5] (darker) and for the ten disulfide-bonded cysteines. (b) OmegaFold and (c) AlphaFold 2 results (alpha carbon RMSD versus average pLDDT for all residues) for the three batches of 50 generated sequences based on this AFP, with the WT sequence included for comparison.

TABLE S1. All protein sequences expressed.

| | Sequence |
|---|---|
| WT AFPIII (GenScript expression evaluation) | MNQASVVANQLIPINTALTLVMMRSEVVTPVGIPAEDIPRLVS MQVNRAVPLGTTLMPDMVKGYAAHHHHHH |
| MPNN1 (GenScript expression evaluation) | MKQPSVVANQLIPINTALTLVMLKAEEVSPQGIPASDIPKLVS KQVTQTIPKGTVIQPEMVKGYVPHHHHHH |
| MPNN2 (GenScript expression evaluation) | MKPASVVANQLIPINTALTLVMLKGEVVSPVGIPYSDIPKLVS KQVVRDIPKGTVIMPEDVKGYVPHHHHHH |
| MPNN3 (GenScript expression evaluation) | MGKKSIVAAQLIPINTALTLVMLGEAEVEPLGIPASDKPKLVS KQVVQTIPKGTVIFENMVKGYQPHHHHHH |
| MPNN4 (GenScript expression evaluation) | MGKKSIVAAQLIPINTALTLVMLGEAVVTPLGIPASDKPKLVS KQTVRTIPKGTVIFENDVKGYKRHHHHHH |
| MPNN5 (GenScript expression evaluation) | MKPKSVVANQLIPINTALTLVMLEAREVEPLGIPASRIPELVS KQVVRDIPKGTVIMPEDVKGYVPHHHHHH |
| MPNN1 (in-house variant expression) | MKQPSVVANQLIPINTALTLVMLKAEEVSPQGIPASDIPKLVS KQVTQTIPKGTVIQPEMVKGYVPLEHHHHHH |
| MPNN2 (in-house variant expression) | MKPASVVANQLIPINTALTLVMLKGEVVSPVGIPYSDIPKLVS KQVVRDIPKGTVIMPEDVKGYVPLEHHHHHH |
| MPNN3 (in-house variant expression) | MGKKSIVAAQLIPINTALTLVMLGEAEVEPLGIPASDKPKLVS KQVVQTIPKGTVIFENMVKGYQPLEHHHHHH |
| MPNN4 (in-house variant expression) | MGKKSIVAAQLIPINTALTLVMLGEAVVTPLGIPASDKPKLVS KQTVRTIPKGTVIFENDVKGYKRLEHHHHHH |
| MPNN5 (in-house variant expression) | MKPKSVVANQLIPINTALTLVMLEAREVEPLGIPASRIPELVS KQVVRDIPKGTVIMPEDVKGYVPLEHHHHHH |
